# Can’t win, don’t try: handling time costs affect foraging choices and vigilance in wild vervet monkeys (*Chlorocebus pygerythrus*)

**DOI:** 10.64898/2026.09.16.752153

**Authors:** Ajay Singh Kang, Wilson Mutebi, Julie A. Teichroeb

## Abstract

Resource handling time is an important cost to foraging animals and this is especially true for low-ranking individuals who may be displaced before they acquire rewards. Handling may also require visual attention that can lead to a trade-off between scanning the environment and focusing on food acquisition. We ran a foraging experiment on wild vervet monkeys where we simultaneously offered them three puzzle boxes in a straight-line array that increased sequentially in handling difficulty and reward value. We found that differences in competitive ability had a large impact on access to long-handling time resources and willingness to invest time in costly problem-solving when social interference was possible. Resources with lower handling time and smaller rewards were accessed by individuals of all dominance ranks but only dominant monkeys that did not fear displacement attempted the high handling time puzzle box. Handling time improved with experience at each puzzle box and contrary to predictions, when vigilance rates were higher, monkeys were also faster at solving puzzle boxes. Thus, there is not necessarily a trade-off between attention to foraging tasks that require handling and vigilance, if a high scanning frequency to both can be balanced with limited scanning duration. These results have important implications for understanding the decision-making of foraging animals that live in competitive groups and the diets of individuals, which may vary with dominance when important resources require long handling times.

**Significance Statement:** Animals are expected to prefer resources that maximize energy intake while minimizing energy expenditure. Many foods require manipulation or processing before they can be consumed, an energetic cost called handling time. Using an experiment on wild, group-living, hierarchical vervet monkeys, we demonstrate that resources with long handling times are especially costly to subordinates, who fear displacement and aggression by dominants. This cost may lead to subordinates not even attempting to access these resources and thus, differing diets for individuals of different rank. Handling that requires visual attention is also suggested to take away from the ability to scan the area for potential dangers, but we show that monkeys could simultaneously invest in high scanning frequencies and shorten their handling times by engaging in many quick glances of short duration.

## Introduction

When deciding whether or not to pursue a resource, animals consider multiple factors, such as quantity, quality, distance, and the levels of predation risk and conspecific competition (MacArthur and Pianka 1966; Galef and Giraldeau 2001; Janson 2007; Sayers and Menzel 2012; Menzel 2012; Teichroeb and Aguado 2016; Kumpan et al. 2019; Arseneau-Robar et al. 2022). According to Optimal Foraging Theory, for a food item to be considered profitable, its net energetic gain must exceed the time and effort required to obtain it (Pyke 1984; Stephens et al. 1986). Consequently, animals are expected to favour foraging options that maximize energetic returns while minimizing costs.

Handling time can be an important foraging cost; this is defined as the time required to manipulate and consume a food item from the moment it is encountered (Cooper and Anderson 2006; Gunst et al. 2010; Sayers and Menzel 2012). Longer handling times may only be advantageous when associated with sufficiently valuable rewards, resulting in greater overall energetic returns. As handling times lengthen, foragers suffer opportunity costs because of reduced time to exploit alternative resources (Brown 1988). In addition, in social species, the costs associated with handling time include the threat of aggressive displacement by dominant individuals, with an associated risk of injury (Janson 1985; di Bitetti and Janson 2001; Arseneau-Robar et al. 2022). Consequently, individuals of different dominance ranks may be expected to adopt alternative foraging decisions based on handling time. Dominant individuals, who are able to exploit high handling time resources with less risk, may be expected to favour profitable resources with high handling times. Whereas subordinates may preferentially select lower-cost alternatives that reduce the likelihood of competitive exclusion and aggression (Janson 1985; Houle et al. 2010; Fowler et al. 2025). Notably, the costs of handling time can be minimized if animals improve their handling speed with experience (Arseneau-Robar *et al*. 2022). Indeed, research has continually shown that prior experience improves animal problem-solving abilities (Birch 1945; Cross et al. 1963; Kolodner and Kolodner 1987; Koton 1988; Cunningham et al. 2011).

Vigilance allows animals to monitor their surroundings and assess potential threats, including both predators and competing conspecifics (Lima and Dill 1990; Beauchamp 2015). In social species, monitoring of conspecifics may be particularly important for lower-ranking individuals, who experience greater vulnerability to displacement and who may need to allocate more attention to dominant group members when accessing resources (e.g., Waite 1987; Gaynor and Cords 2012). A trade-off may also exist for food items that require visual attention while handling because attention given to food extraction theoretically decreases the vigilance that can be allocated to the surrounding environment (i.e., predators and conspecifics) (Kaby and Lind 2003; Cowlishaw et al. 2004; Baker et al. 2011; Teichroeb and Sicotte 2012). The degree to which individuals adjust vigilance in response to the presence, proximity, and identity of conspecifics may therefore influence their ability to optimize resource acquisition.

One way to ascertain how behaviour and decision-making are influenced by trade-offs like visual attention to food vs. competitors is to use experiments that allow subjects to evaluate the costs and benefits of foraging opportunities within social environments. In the wild, these types of experiments are particularly relevant because animals are tested under the full suite of evolutionary constraints that they experience daily (Janson 2012). In this study, we presented a multi-destination foraging array to wild vervet monkeys at Nabugabo, Uganda consisting of three platforms in a row 5 m apart, each equipped with a foraging puzzle box. The three puzzle boxes, designated as Easy, Medium, and Hard, required monkeys to retrieve rewards using increasingly complex techniques and differed in reward value. The Easy box required a no-manipulation reach-in, the Medium box, a pulling technique with a reach-in, and the Hard box, a twisting-and-pulling combination with a reach-in, and reward value was higher with increasing complexity (Fig. 1). Unlike many captive studies, the monkeys received no prior training and were free to manipulate and investigate each puzzle to discover the solution independently in the weeks prior to the boxes being placed in the array. This experimental design allowed individuals to first learn the box solutions and then choose among foraging options where they had knowledge of the variation in the costs of reward acquisition.

**Figure 1.**
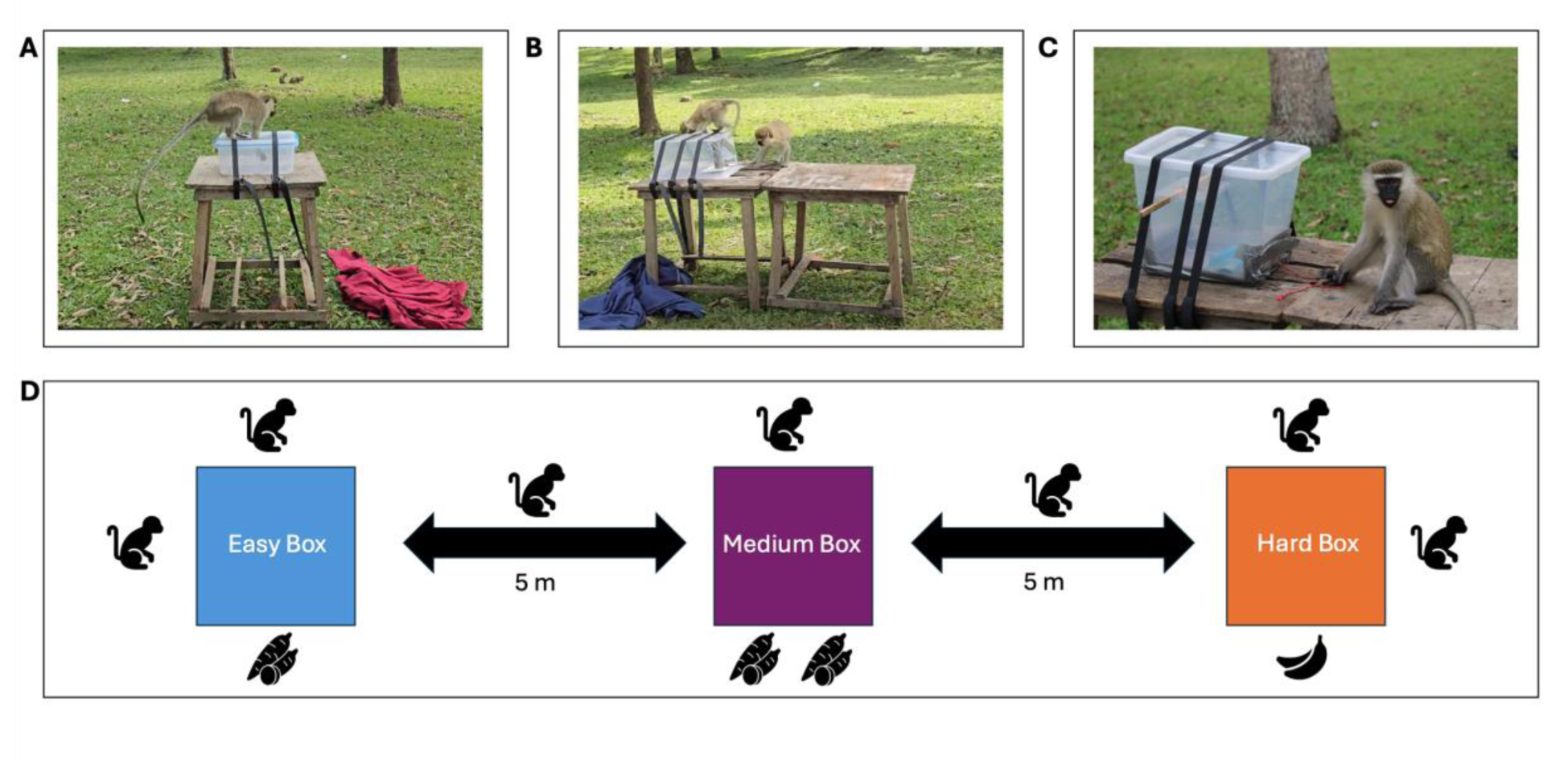
Experimental apparatus and task difficulty progression. The multi-destination foraging array used to assess problem-solving behaviour in wild vervet monkeys (*Chlorocebus pygerythrus*) at Nabugabo, Uganda. Panels (A–C) depict individuals interacting with the puzzle box apparatus across the three experimental conditions: (A) an individual extracting the reward from the Easy box, which was solved by reaching into an opening to retrieve the food reward; (B) an individual manipulating the Medium box, which required pulling a Styrofoam slab using an attached string to access the reward; and (C) an individual following the completion of the Hard box condition, which required a combination of twisting and pulling actions to retrieve the reward. Panel (D) provides a schematic representation of the multi-destination array that the puzzle boxes were placed in, illustrating variation in reward placement and the increasing levels of task difficulty across conditions. For the final phase of the experiment, the Easy box was baited with a chunk of cassava, the Medium box had a larger piece of cassava, and the Hard box was baited with a piece of more preferred banana.

Multi-destination foraging arrays have been used to investigate the cognitive abilities and decision-making of wild vervet monkeys at Lake Nabugabo, Uganda, for over a decade (Teichroeb and Chapman 2014; Teichroeb and Aguado 2016; Teichroeb and Smeltzer 2018; Kumpan et al. 2019; Arseneau-Robar et al. 2022, 2023; Fowler et al. 2025). Vervet monkeys are matrilineal, primarily frugivorous, semiterrestrial cercopithecines that live in stable social groups characterized by linear dominance hierarchies (Teichroeb 2015; Cancelliere et al. 2018; Fowler et al. 2025). However, these hierarchies do not always rigidly determine access to resources, as recent social interactions and affiliative relationships can promote tolerance between conspecifics (Borgeaud and Bshary 2015; Fowler et al. 2025). This dynamic social system provides an ideal study system for investigating foraging decision-making.

Based on our study design, we hypothesized that the cost in handling time needed to solve each puzzle box and the associated rewards would influence participation by vervets. In particular, as difficulty increased and handling time grew, the threat of displacement by dominants would increase. The Hard box with its high handling time and better reward was predicted to be especially attractive to dominant monkeys, who also had the ability to monopolize it. Thus, we predicted that the Easy box would have a greater proportion of participation by lower-ranking monkeys, relative to the Medium box, and the Hard box would have the lowest number of attempts by subordinates. When monkeys had the opportunity to navigate through the whole array, we predicted that all monkeys would choose to go in decreasing order of difficulty (Hard, Medium, Easy) because the best reward was in the Hard box and they would want to secure it first before competitors could arrive. For all boxes, regardless of difficulty, we hypothesized that prior experience would improve handling times (Arseneau-Robar et al. 2022, 2023) and that handling time would be faster when audience member of the same or higher-ranking dominance tier were closer rather than further away, because the heightened pressure of conspecific competition would spur individuals to solve the box as fast as possible. Vigilance was hypothesized to be greater for subordinate individuals relative to more dominant monkeys at every box. Note that though animal vigilance has been defined variously by its intensity and its target (Blanchard and Fritz 2007; Beauchamp 2015), in the case of our foraging experiments, vigilance was almost always social vigilance induced by conspecific competitors. Vigilance rates were predicted to be highest when same or higher-ranking individuals were nearby and lowest when they were more distant, particularly while subordinate individuals were trying to solve puzzle boxes with long handling times. If a trade-off was present between attention to the puzzle boxes and vigilance, we predicted slower handling times when vervets were more vigilant to competitors.

## Methods

### Study Site & Subjects

This study took place at Lake Nabugabo, situated in the Masaka District of central Uganda (0°22’12“S, 31°54’0”E). Lake Nabugabo is a satellite lake of Lake Victoria and spans an area of 8.2 by 5 kilometers and lies at an altitude of 1136 meters. The subjects of this study are a well-habituated group of vervet monkeys (*Chlorocebus pygerythrus*) known as KS group, which has been under continuous observation since 2016 (Li et al. 2021). The home range of KS group encompasses a diverse habitat mosaic, including open woodland, grassland, farmland, and a limited number of buildings. The landscape surrounding Lake Nabugabo is significantly influenced by human activity. While the vervet monkeys at the study site primarily rely on natural food sources, their diet is supplemented by anthropogenic foods, including crops, discarded human foods, and tourist-provided resources (Chapman et al. 2016). This human-derived food remains finite however, and insufficient to fully satiate all members of the group, leading to prevalent feeding competition. At the time of this study, KS group included 29-31 individuals, 2 adult males, 8 adult females, 1 subadult male, 3 subadult females, and 15-17 juveniles/infants, all of which were individually identified based on their unique physical characteristics. Dominance status was assessed by recording aggressive behaviours, including displacements, and submissive responses between individuals whenever these were observed. However, due to some rank instability during the period of the experiment, we cautiously assigned individuals to dominance tiers rather than assigning them an ordinal rank. Dominance was classified in tiers 1-4 (with Tier 1 being the most dominant individuals and Tier 4 being the most subordinate).

### Data Collection

Behavioural data were collected via video recordings and all occurrence sampling during field experiments. Data collection periods occurred six days a week between 08:00 and 16:00 for a duration of three months (June-August 2025). During experimental trials, the monkeys were provided access to a multi-destination array comprised of platforms baited with puzzle boxes that varied in difficulty and rewards. All boxes were secured to the platforms with tension cables to prevent monkeys from twisting, turning, or removing the boxes during manipulation (Fig. 1). The Easy box consisted of a clear plastic box with a hole positioned on one side of a transparent lid, allowing monkeys to access a small piece of cassava by inserting their arm through the opening (One step - reach in; Fig. 1A). The Medium box consisted of a clear plastic box that was larger than the Easy box and contained a white Styrofoam slab attached to a string. Monkeys were required to pull the string toward themselves, bringing the slab within reach to retrieve a large piece of cassava (Two steps – pull string, reach in; Fig. 1B). Lastly, the Hard box consisted of a plastic box larger than both the Easy and Medium boxes and required monkeys to perform a two-step manipulation sequence to retrieve a banana wedge. A small, rectangular plastic plate was attached to the centre of a rectangular dowel rod positioned across the upper interior of the box. Beneath the dowel was a Styrofoam slab attached to a string, like the setup used in the Medium box. To retrieve the reward, monkeys first had to twist the dowel to rotate the plate and release the reward onto the slab, then pull the string toward themselves to bring the slab within reach and extract the reward from the box (Three steps – flip plate, pull string, reach in; Fig. 1C).

Prior to the presentation of the multi-destination array, a time of one week was allocated for each puzzle box to be learned by the monkeys (three weeks in total). Each box was presented individually for the full week or until 20 out of the 29 individuals successfully solved the box and obtained a reward. Following these three weeks, the boxes were placed on platforms in a straight line 5 m from one another. For three weeks, each box was baited with an equal amount of food (e.g., banana and/or cassava). Following this, until the end of the experiment (four weeks) the boxes were baited with varying reward levels such that the Easy and Medium boxes contained cassava (with the Medium box containing a larger volume of the reward) and the Hard box contained a chunk of banana (Fig. 1). Cassava is higher in starch and lower in sugar relative to banana, so the latter was preferred by the monkeys (Montagnac et al. 2009; Phillips et al. 2021). Boxes were baited and then covered with a drop cloth to discourage the vervets from trying to access the rewards before trials began. Removing the cloth(s) acted as a signal to the monkeys that they could approach, make a box selection within the array, and begin to try to retrieve rewards.

Video recordings were used to code and analyze the behaviours of the subjects during experiments. It was not possible to record data blind because our study involved focal animals in the field. One observer (ASK) video recorded the solver, while an assistant (WM) recorded the audience around the platform(s) on a data sheet. During experimental trials where the boxes were in the array, each box was recorded, for later coding, using a camera attached to a tripod, except for the Easy box which was recorded with the observer holding the camera. During each experimental trial the identity of the monkey obtaining the reward, the composition of the audience and their distance from the experiments (in bins: 0-10 m, 10-20 m, 20-30 m, 30-40 m, 40-50 m), the order of platform visitation, and any social interactions were recorded for analysis. Video recordings were analyzed later for precise measurements of handling time (i.e., time from touching the box to acquiring the reward) and vigilance rate (i.e., the total number of head turns of at least 45° per second by the focal individual). These variables were coded into Microsoft Excel for each trial by a single observer (ASK) to maintain internal consistency in coding.

## Data Analyses

All statistical analyses were conducted in Python (version 3.13.9; Python Software Foundation) using the packages *pandas*, *numpy*, *scipy*, *statsmodels*, *matplotlib*, and *seaborn*. Data processing, cleaning, and visualization were performed using *pandas*, *numpy*, *matplotlib*, and *seaborn*, while statistical modelling and hypothesis testing were conducted using *scipy* and *statsmodels* (Seabold and Perktold 2010; Virtanen et al. 2020). Only successful and complete trials were included in the analyses. Trials were excluded if the apparatus was dislodged from the platform or removed from the table, as these events prevented individuals from completing the intended problem-solving sequence. Additionally, trials involving cooperative manipulation, where multiple individuals contributed to reward retrieval (e.g., for the Hard box, one individual twisting the rod while another pulled the slab), were excluded.

Because individuals contributed multiple trials throughout the experiment, repeated observations from the same monkey were not considered independent. Therefore, all analyses involving continuous behavioural measures were conducted using linear mixed-effects models (LMMs), with individual identity (ID_1) included as a random effect to account for individual differences in baseline problem-solving ability and repeated measures within individuals (Zuur et al. 2009). Models were fitted using restricted maximum likelihood estimation. Continuous time-based measures were log-transformed using a log(x+1) transformation to reduce the influence of positive skew in latency measures. Model assumptions were evaluated through inspection of residual distributions and convergence diagnostics. Predictor variables were examined for potential multicollinearity prior to model interpretation. Statistical significance was assessed using an alpha level of 0.05, and model estimates were interpreted alongside effect sizes and confidence intervals.

### Initial Performance Across Puzzle Boxes

To compare problem-solving performance among puzzle box conditions and ensure that our design did indeed lead to increases in difficulty for vervets with each introduced box, analyses were conducted using only individuals who completed all three experimental conditions. Restricting analyses to these individuals allowed direct within-subject comparisons of performance across Easy, Medium, and Hard boxes while minimizing variation caused by differences in participation among individuals. First-trial performance was extracted for each individual in each condition and these first-trial handling times were compared using LMMs between conditions (Easy vs. Medium, Easy vs. Hard, Medium vs. Hard) with individual identity included as a random effect. Because these data consisted of matched observations from the same individuals across three conditions, a Friedman test was additionally conducted as a non-parametric within-subject comparison. Kendall’s *W* was calculated as an effect size measure for the Friedman test to quantify the magnitude of differences in first-trial performance among puzzle box conditions.

### Puzzle Box Participation

To determine whether dominance status influenced participation in puzzle box conditions when all three apparatuses were simultaneously available, we examined whether individuals from different dominance tiers differed in their completion of each puzzle box condition. Because individuals could interact with and successfully complete more than one puzzle box across experimental trials, participation was defined independently for each condition based on whether an individual completed at least one successful trial for the Easy, Medium, or Hard box. For each dominance tier, we calculated the number and proportion of individuals that completed each puzzle box condition. To statistically assess whether dominance status predicted participation, we conducted a binary logistic regression analysis for the Hard box with completion status (completed/not completed) as the response variable and dominance tier as the predictor. Because all individuals completed the Easy box and most completed the Medium box, these conditions lacked variation in completion and could not be analysed statistically with the logistic regression. Thus, to further assess whether dominance status and puzzle box difficulty jointly influenced participation across all three conditions, we conducted a binomial generalized linear mixed-effects model (GLMM). Completion status was used as the response variable, dominance tier and puzzle box condition were the fixed effects, and individual identity was included as a random intercept. Puzzle box difficulty was examined as an ordinal predictor (Easy = 1, Medium = 2, and Hard = 3). Dominance tier was also used as an ordinal predictor, with lower numerical values representing higher-ranking individuals (Tier 1) and higher numerical values representing lower-ranking individuals (Tier 4). Odds ratios were calculated to quantify the change in probability of completing the Hard box with increasing dominance tier.

### Path Choice Analysis

Behavioural strategy differences in the order of platform visitation were examined using chi-square tests of independence and permutation-based chi-square tests. These analyses could only be performed on SOY and TBR (the alpha female and alpha male), as these were the only individuals that completed at least three trials in which they progressed through the entire array. Permutation tests were conducted because behavioural strategies were not evenly distributed across individuals and some categories contained relatively few observations. Effect sizes were quantified using Cramér’s V to estimate the strength of associations between individual identity and strategy use.

### Effect of Experience, Vigilance, and Audience Distance on Handling Time

To first examine whether handling time improved with experience, we calculated the percent reduction in handling time across all trials in each condition. We only included data from individuals that had a minimum of five trials for that particular box. Then, to assess and control for multiple variables that may have impacted handling time, we ran a LMM for each trial condition (Easy, Medium, and Hard) that assessed log-transformed handling time as the dependent variable and included trial number for each individual as a measure of experience, vigilance rate during that trial, and audience distance (as a categorical variable because this was collected in bins) as fixed effects. Individual ID was included in each model as a random effect to control for repeated sampling of the same individuals over time. We log-transformed handling time because these data were positively skewed and the transformation improved the normality and homoscedasticity of the residual, thereby better meeting the assumptions of the LMM.

### Impacts of Dominance Rank and Audience Distance on Vigilance

Vigilance behaviour was analysed to determine whether it was affected by dominance tier or the proximity of same-or higher-ranking individuals while monkeys completed the experiment. Again, separate LMMs were conducted for each puzzle box condition. Audience distance was included as a categorical fixed effect and dominance tier was included as a continuous fixed effect, with individual ID included as a random effect.

## Results

### Initial Performance Across Puzzle Boxes

We first assessed whether puzzle box difficulty increased as intended by examining first-trial handling times among individuals that completed all three experimental conditions (N = 5 individuals that completed each condition, N = 15 first-trial observations; Fig. 2). A mixed-effects model revealed that first-trial handling time was significantly longer in the Hard condition compared with the Easy condition (LMM: β = 1.223 ± 0.426 SE, *P* = 0.004). Performance on the Medium box did not differ significantly from the Easy box (β = 0.187 ± 0.426 SE, *P* = 0.661). However, a post-hoc contrast revealed that handling time was significantly longer for the Hard box compared with the Medium box (β = 1.036 ± 0.426 SE, *P* = 0.015). A Friedman test comparing the three matched conditions did not reach statistical significance (χ² = 5.20, *P* = 0.074), but showed a trend, and the effect size indicated a large effect of puzzle box condition (Kendall’s *W* = 0.52), suggesting that task difficulty contributed to meaningful differences in initial problem-solving performance despite the limited sample size of first attempts by individuals.

**Figure 2.**
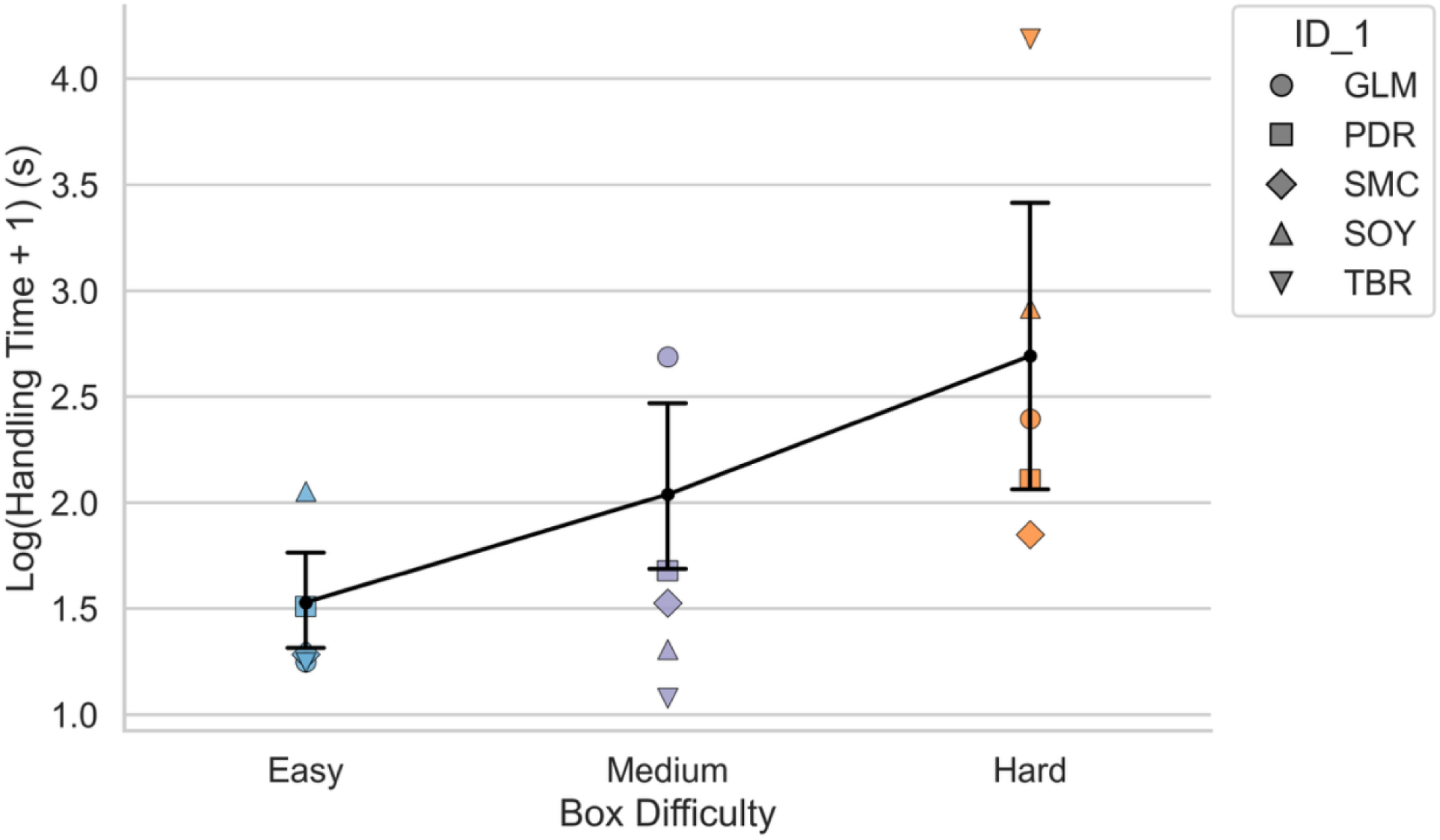
Handling time across Easy, Medium, and Hard puzzle box conditions for first attempts by individual vervets (*Chlorocebus pygerythrus*) that completed all three experimental conditions at Nabugabo, Uganda. Points represent individual first-trial performances, with different shapes indicating individual identities and black points and lines represent the mean handling time across conditions, with error bars showing ± standard error.

### Puzzle Box Participation

Across all experimental trials, individuals contributed different numbers of observations for the three puzzle box conditions, reflecting variation in participation and engagement with the apparatus. The Easy box received the highest level of participation across all dominance ranks (Tier 1: 3/3, 100%; Tier 2: 3/3, 100%; Tier 3: 4/4, 100%; Tier 4: 7/7, 100%) (Table 1). For the Medium box, most individuals across all dominance tiers completed the task, although participation was lower among Tier 4 individuals (Tier 1: 3/3, 100%; Tier 2: 3/3, 100%; Tier 3: 4/4, 100%; Tier 4: 5/7, 71.4%). In contrast, completion of the Hard box differed substantially across dominance tiers, with all Tier 1 individuals completing the condition (3/3, 100%), compared with fewer Tier 2 individuals (2/3, 66.7%) and no Tier 3 or Tier 4 individuals (Tier 3: 0/4, 0%; Tier 4: 0/7, 0%) (Table 1). A binary logistic regression examining whether dominance predicted Hard box completion revealed a significant negative relationship between dominance tier and completion probability (β =-1.661, SE = 0.787, *P* = 0.035), with lower dominance rank being associated with reduced odds of completing the Hard box (odds ratio = 0.190).

**Table 1.** Participation in each puzzle-box condition by vervets (*Chlorocebus pygerythrus*) of different dominance tiers at Lake Nabugabo, Uganda. The proportion of individuals within each dominance tier that successfully completed at least one Easy, Medium, or Hard puzzle-box trial while all three puzzle boxes were simultaneously available. Percentages are calculated relative to the total number of individuals within each dominance tier.

| Dominance Tier | Individuals in Tier (N) | Easy Box (%) | Medium Box (%) | Hard Box (%) |
| --- | --- | --- | --- | --- |
| 1 | 3 | 100.0 | 100.0 | 100.0 |
| 2 | 3 | 100.0 | 100.0 | 100.0 |
| 3 | 4 | 100.0 | 100.0 | 0.0 |
| 4 | 7 | 100.0 | 71.4 | 0.0 |

To assess whether participation changed systematically with increasing puzzle-box difficulty, a binomial GLMM treated difficulty as an ordered predictor. The model indicated a negative effect of dominance tier on completion probability (β =-1.266, 95% CrI = [-2.510, - 0.079]), with lower-ranking individuals showing reduced odds of completing the puzzle boxes. The odds ratio associated with dominance tier (0.283, 95% CrI = [0.081, 0.924]) indicated that with each increase in tier there was approximately a 72% reduction in the odds of box completion. The effects of increasing puzzle box difficulty were negative but did not receive clear statistical support (β =-1.199, 95% CrI = [-2.826,-0.346]; OR =0.307, 95% CrI = [0.059, 1,414]). Similarly, the difficulty x dominance tier interaction did not clearly differ from zero (β =-0.424, 95% CrI = [-0.873, 0.001]; OR = 0.656, 95% CrI = [0.418, 1.001]).

### Path Choice

Individuals exhibited non-random patterns in the order in which they completed the puzzle boxes during trials where all three boxes were completed by the same focal individual (N = 2) (Fig. 3). For SOY, the alpha female of the KS group (N = 33), the distribution of approach orders differed significantly from random expectations (χ² = 18.22, df = 6, *P* = 0.006). This result was confirmed using a permutation-based chi-square test (permutation *P* = 0.0015). The strength of this association was large (Cramér’s V = 0.53), indicating that SOY had a consistent pattern in the sequence in which she approached the puzzle boxes. Similarly, TBR, the alpha male of the KS group (N = 24), also demonstrated a non-random distribution of approach orders (χ² = 10.05, df = 4, P = 0.040). This result was supported by the permutation-based chi-square test (permutation *P* = 0.0339), with a medium-to-large effect size (Cramér’s V = 0.46), showing that TBR also showed a consistent preference for certain approach sequences. Although both individuals demonstrated consistent approach strategies, their preferred sequences differed. SOY most frequently approached the boxes in an Easy–Medium–Hard order, whereas TBR more frequently initiated trials by approaching the Hard box.

**Figure 3.**
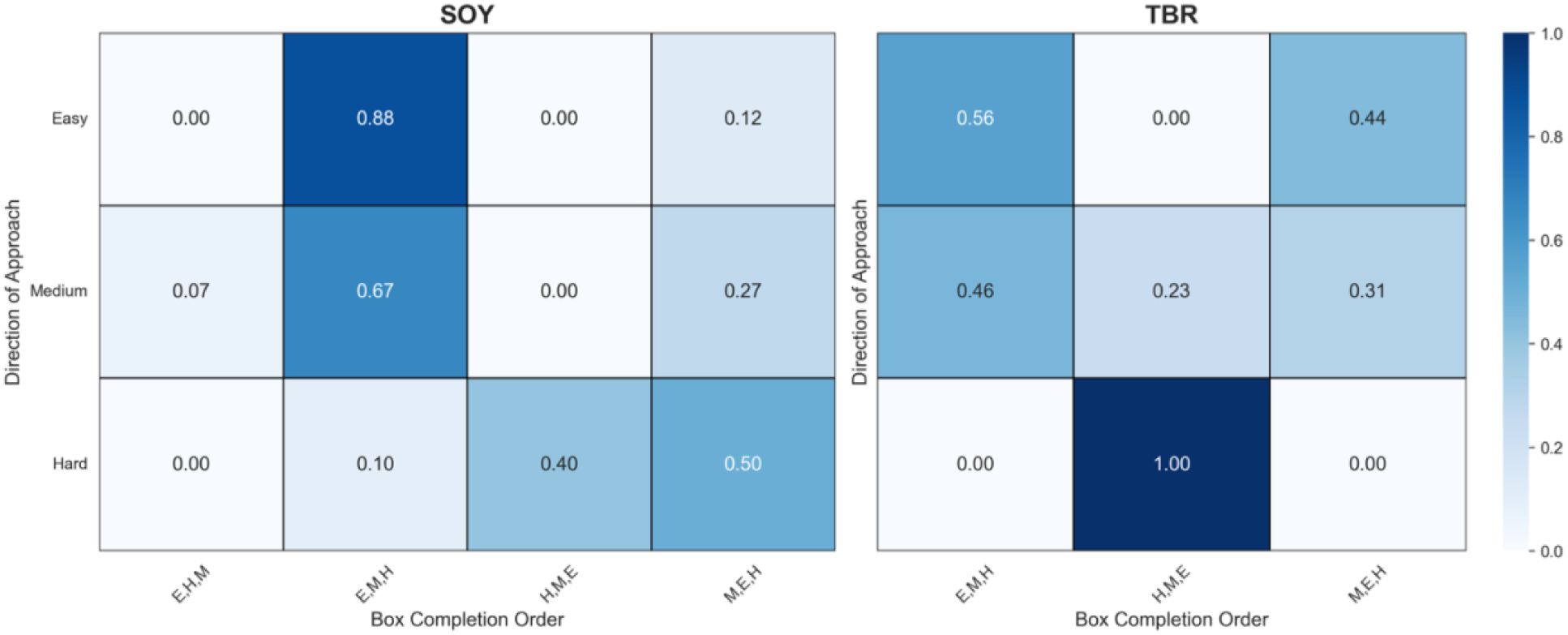
Proportion of box completion orders by direction of approach (DOA) for SOY and TBR, the alpha female and alpha male of vervet KS Group at Nabugabo, Uganda. Rows indicate the initial direction of approach (Easy, Medium, Hard), and columns indicate the order in which the three puzzle boxes were completed.

### Effect of Experience, Vigilance, and Audience Distance on Handling Time

Across all three puzzle box conditions, monkeys decreased handling times with repeated exposure to the apparatus. In the Easy box condition (N = 21 individuals; minimum of five trials per individual), individuals showed a 27% reduction in handling time across 310 trials. Similarly, monkeys in the Medium box condition (N = 17 individuals; minimum of five trials per individual) improved by 32.2% across 389 trials. In the Hard box condition, five individuals who completed a minimum of five trials reduced their handling time by 31% across 377 trials.

LMMs demonstrated that, for the Easy box, handling time significantly decreased with experience (β =-0.001 ± 0.000 SE, *P* < 0.001) and was strongly associated with vigilance rate, with individuals exhibiting higher rates of head turns completing trials more quickly (β =-0.651 ± 0.024 SE, *P* < 0.001). However, the distance to audience members of the same or higher dominance tier did not significantly impact handling times for this box (Table 2). Similarly, handling times at the Medium box decreased with both experience (β =-0.001 ± 0.000 SE, *P* < 0.001) and with increased vigilance rate (β =-0.799 ± 0.027 SE, *P* < 0.001). Additionally shorter handling times were seen at this box when audience members of the same or higher dominance tier were 20-30 m away (LMM: β =-0.040 ± 0.019 SE, *P* = 0.037) but these were longer when these audience members were 30-40 m away (LMM: β = 0.125 ± 0.037 SE, *P* = 0.001). Likewise, for the Hard box, vigilance rate had the strongest impact on performance, with higher vigilance rates associated with substantially shorter handling times (LMM: β =-3.134 ± 0.103 SE, *P* < 0.001) and experience also led to shorter handling times (LMM: β =-0.00047 ± 0.000 SE, *P* < 0.001). Distance to audience members of the same or higher dominance tier was not a significant predictor of handling time for the Hard box (Table 2).

**Table 2.**
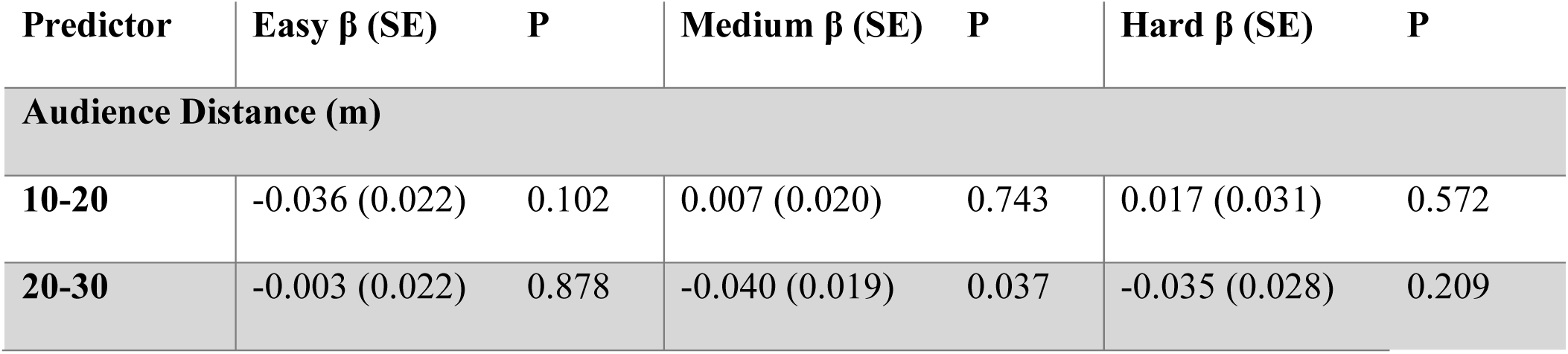
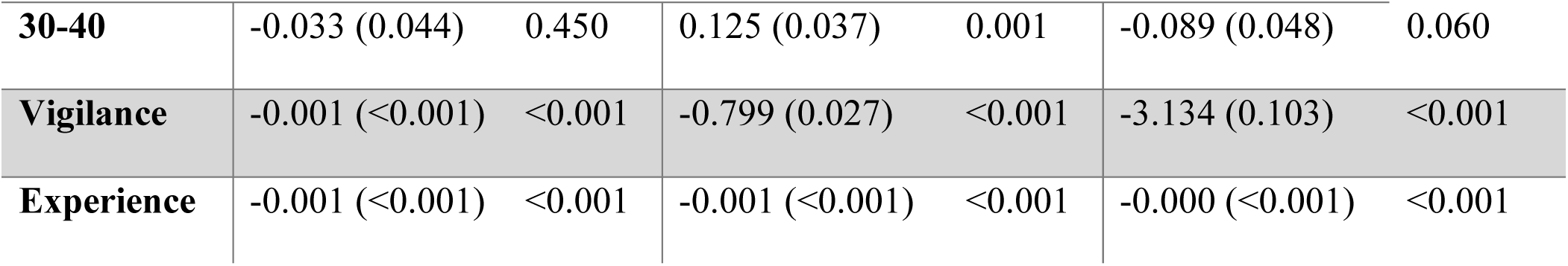
Results of Linear Mixed Models examining the effects of audience distance, vigilance, and experience on handling time across puzzle box conditions for vervet monkeys (*Chlorocebus pygerythrus*) at Nabugabo, Uganda.

| Predictor | Easy $\beta$ (SE) | P | Medium $\beta$ (SE) | P | Hard $\beta$ (SE) | P |
| --- | --- | --- | --- | --- | --- | --- |
| **Audience Distance (m)** |  |  |  |  |  |  |
| **10-20** | -0.036 (0.022) | 0.102 | 0.007 (0.020) | 0.743 | 0.017 (0.031) | 0.572 |
| **20-30** | -0.003 (0.022) | 0.878 | -0.040 (0.019) | 0.037 | -0.035 (0.028) | 0.209 |
| **30-40** | -0.033 (0.044) | 0.450 | 0.125 (0.037) | 0.001 | -0.089 (0.048) | 0.060 |
| **Vigilance** | -0.001 (<0.001) | <0.001 | -0.799 (0.027) | <0.001 | -3.134 (0.103) | <0.001 |
| **Experience** | -0.001 (<0.001) | <0.001 | -0.001 (<0.001) | <0.001 | -0.000 (<0.001) | <0.001 |

### Impacts of Dominance Rank and Audience Distance on Vigilance

Vigilance behaviour was examined relative to an individual’s dominance tier and the distance of audience members of the same or higher dominance tier (Fig.4). For the Easy box, dominance tier did not significantly influence vigilance rates (β =-0.033 ± 0.051 SE, *P* = 0.524) but individuals exhibited fewer vigilant head turns when same or higher-ranking individuals were positioned 20–30 m away (β =-0.089 ± 0.042 SE, *P* = 0.035), while other distance categories did not differ significantly (Table 3; Fig. 4A). Vigilance rate at the Medium box was also not influenced by dominance tier (β =-0.004 ± 0.036 SE, *P* = 0.910) but was found to be significantly higher when audience members were at intermediate and far distances (20–30 m: β = 0.092 ± 0.030 SE, *P* = 0.002; 30–40 m: β = 0.188 ± 0.058 SE, *P* = 0.001) (Fig. 4B). Lastly, at the Hard box, dominance tier did significantly influence vigilance rate, with higher tier numbers (lower-ranking individuals) associated with higher vigilance rates (β = 0.177 ± 0.081 SE, *P* = 0.029), and though individuals tended to make fewer vigilant head turns when audience members were farther away, but the effect was small and not statistically significant (Table 3; Fig. 4C).

**Figure 4.**
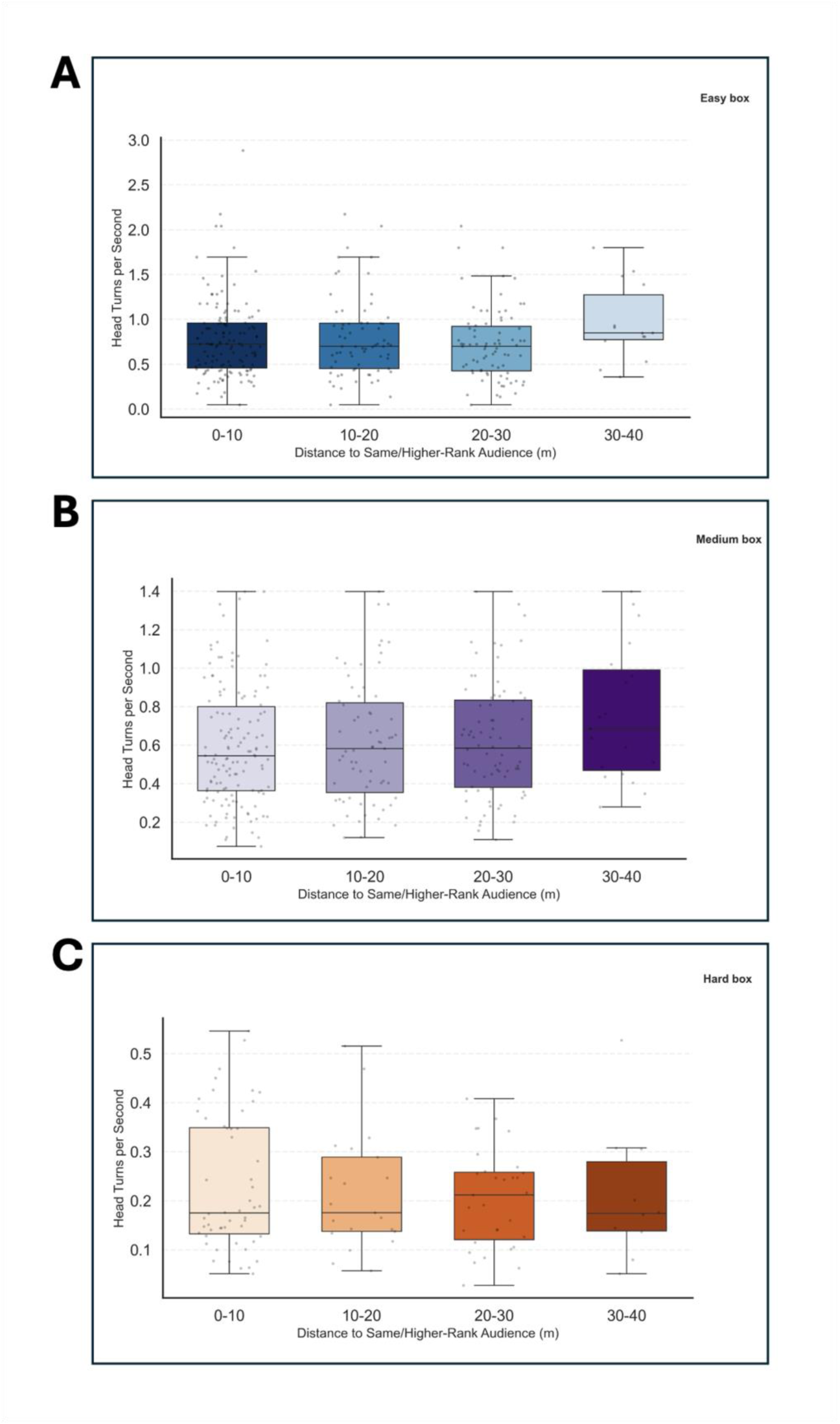
Vigilance rate (Head turns per second) for vervet monkeys (*Chlorocebus pygerythrus*) at Nabugabo, Uganda when the distance to the nearest individual of same or higher rank was within 10 m, 10-20 m, 20-30 m, and 30-40 m, for individuals, that completed the (A) Easy box, (B) Medium box, and (C) Hard box.

**Table 3.** Results of Linear Mixed Effects Models examining the effects of audience distance and dominance tier on vigilance rate across puzzle box conditions for vervet monkeys (*Chlorocebus pygerythrus*) at Nabugabo, Uganda.

| Predictor | Easy $\beta$ (SE) | P | Medium $\beta$ (SE) | P | Hard $\beta$ (SE) | P |
| --- | --- | --- | --- | --- | --- | --- |
| **Audience Distance (m)** |  |  |  |  |  |  |
| **10-20** | -0.005 (0.044) | 0.904 | 0.006 (0.033) | 0.857 | -0.020 (0.027) | 0.452 |
| **20-30** | -0.089(0.042) | 0.035 | 0.092 (0.030) | 0.002 | -0.034 (0.024) | 0.153 |
| **30-40** | -0.120 (0.089) | 0.177 | 0.188 (0.058) | 0.001 | -0.021 (0.035) | 0.544 |
| **Dominance Tier** | -0.033 (0.051) | 0.524 | -0.004 (0.036) | 0.910 | 0.177 (0.081) | 0.029 |

## Discussion

This study examined how increasing handling time costs for a set of resources impacted vervet monkey foraging decisions and vigilance in a wild group. Our initial analyses assessed whether the handling time of first attempts, by monkeys that completed each box, corroborated our goal of designing an experiment with increasing puzzle box difficulty. We did indeed find that handling times were longer for monkeys as they progressed from the Easy (one step sequence), to the Medium (two-step sequence), to the Hard box (three-step sequence). Overall, our results demonstrate that handling time cost and the resulting reward strongly influenced which individuals chose each box, with dominance rank as a key determining factor. As expected, individual experience improved performance, with handling times decreasing substantially across repeated trials, regardless of puzzle box difficulty. Against expectations, vigilance was strongly positively associated with handling time with individuals completing each box more quickly when they made more vigilant head turns. Only the Hard box, with the highest handling time, showed significantly greater vigilance for lower-ranking individuals relative to more dominant monkeys, while for the other boxes, although vigilance was higher for subordinates this was not significant (Fig. 5). The distance of same or higher ranked audience members did not show clear effects on either handling time or vigilance, but patterns did not support our predictions that closer competitors would lead to shorter handling times and greater vigilance.

**Figure 5.**
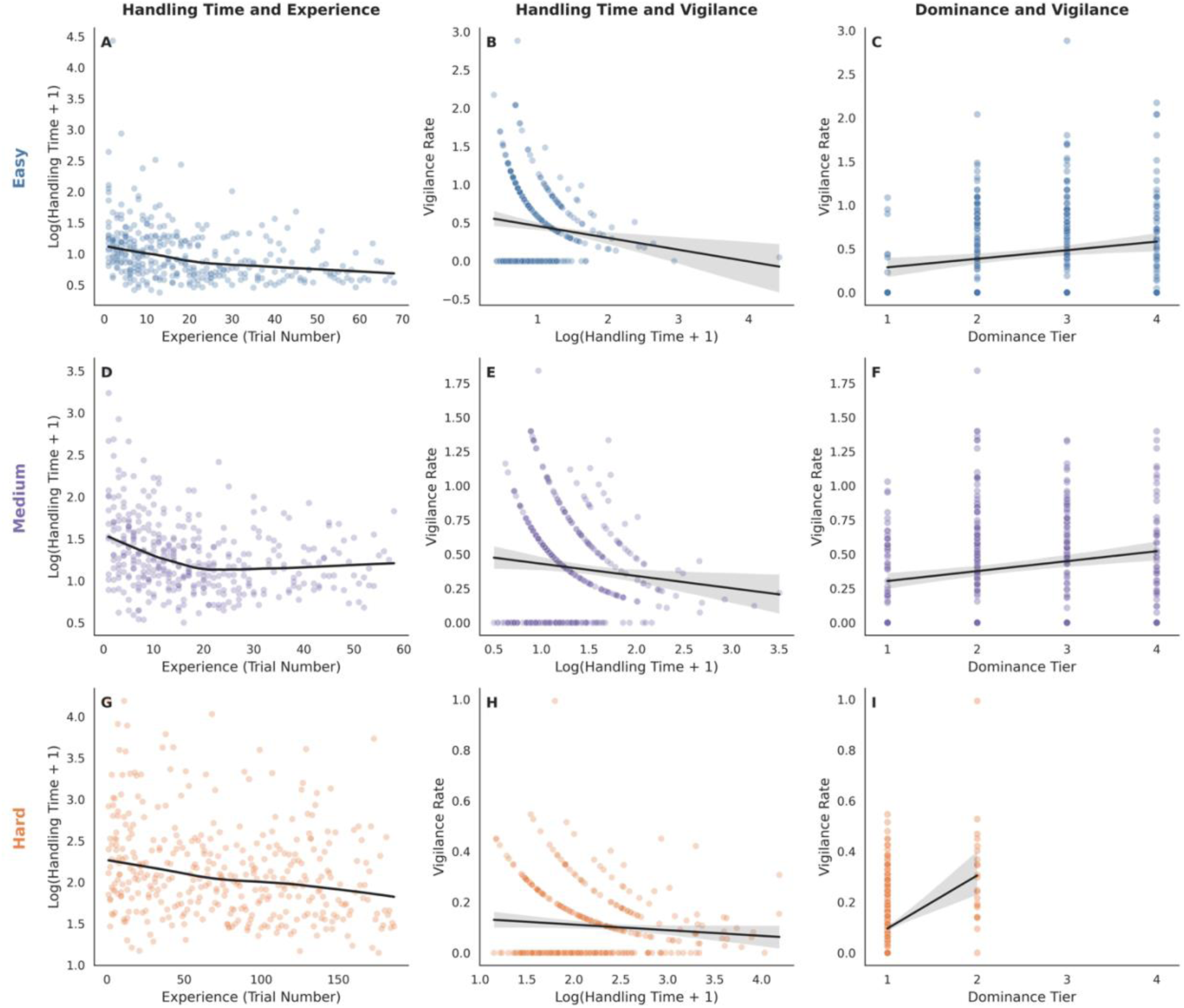
Relationships between task experience, handling time, vigilance rate, and dominance tier across puzzle box difficulties for vervet monkeys (*Chlorocebus pygerythrus*) at Nabugabo, Uganda. Rows represent the difficulty of puzzle box conditions: Easy (top row; A-C), Medium (middle row; D-F), and Hard (bottom row; G-I). Columns represent behavioural relationships: Log-transformed handling time plotted against trial numbers (A, D, G), Vigilance rate plotted against log-transformed handling time (B, E, H), and Vigilance rate across dominance rank tiers (1-4) (C, F, I).

The distribution of participation across puzzle box conditions provides evidence that task difficulty, reward value, and dominance status interacted to influence engagement with the apparatus. As predicted, the Easy box was accessible to individuals across all dominance ranks and all individuals that participated in the experiment completed this condition. The low handling time cost and reduced manipulation demands of the Easy box allowed individuals across the hierarchy to engage with the task without substantial social or cognitive constraints. The Medium box had slightly lower participation by the lowest-ranking monkeys, while participation for the Hard box was strongly associated with dominance status, with only Tier 1 and Tier 2 individuals successfully completing this condition. The negative relationship between dominance tier and completion of the boxes with longer handling times indicates that, as predicted, the threat of displacement, before even getting the reward, discouraged lower-ranking individuals from attempting the harder boxes. Dominant individuals, who had little risk of displacement, were freer to spend time at the Hard box trying to solve it. The better reward in this box in the final set of trials made it even more attractive to high-ranking individuals.

Thus, differences in competitive ability had a large impact on access to long-handling time resources and willingness to invest in costly problem-solving when social interference was possible. This has important implications for how we understand diets in wild animal groups, particularly when dominance hierarchies are strict and resources have long handling times. In species experiencing these conditions, it would be more accurate to study diet individually, relative to dominance rank, rather than across a group as a whole, as is typically done. Studies in other species have also shown rank-based differences in access to valuable resources that lead to differing diets for animals of different ranks (e.g., *Canis lupis*, Atwood and Gese 2008; *Cercopithecus mitis,* Foerster et al. 2011; *Rhinopithecus roxellana*, Guo et al. 2020; *Pan troglodytes*, Murray et al. 2006; *Gorilla beringei*, Wright et al. 2014) and our work shows that long handling times for rare resources exacerbate this effect.

Although we predicted that monkeys that navigated through the whole array would go in decreasing order of puzzle box difficulty, in practice, we were not able to test this hypothesis. Likely owing to the low levels of participation by lower-ranking monkeys at the Hard box, we only had two individuals that navigated through the array as a whole, and these were the two most dominant animals in the group. The alpha male (TBR) and the alpha female (SOY) were able to move through the whole array on occasion, but each showed different preferred routes. TBR started most frequently with the hard box, which is what we predicted monkeys would be most likely to do - so that they could ensure they got the best reward before competitors. However, SOY showed the opposite pattern, most often completing the array by going Easy-Medium-Hard. Thus, the question of how variability in handling time costs may influence the order of food selection needs to be addressed in subsequent studies.

In the literature, it is commonly found that prior experience improves problem-solving performance in animals (e.g., Cross et al. 1963; Visalberghi and Limongelli 1994; Cunningham et al. 2011; Ebel and Call 2018; Arseneau-Robar et al. 2022, 2023) and our results supported our second hypothesis that this would be the case for vervets in our experiment. Across all puzzle box conditions, monkeys reduced their handling times with repeated exposure, demonstrating that individuals learned from their own interactions with the apparatus. However, the rate of improvement differed according to task difficulty. Learning trajectories for the Easy and Medium puzzle boxes were similar, with handling times declining rapidly before approaching a plateau. In contrast, the Hard puzzle box exhibited a slower learning trajectory (Fig. 5), indicating either that: 1) increased complexity reduced the rate at which individuals acquired efficient solutions; or 2) that the higher ranked individuals that completed the Hard box were less motivated to learn. A previous experiment with this study group also showed improved handling time with experience on an apparatus similar to the Easy box but demonstrated that subordinates showed much quicker learning trajectories than dominants (Arseneau-Robar et al. 2023). It would have been informative to perform a similar analysis on the Hard box in the present study, but the long handling time and increased reward was such that high-ranking individuals dominated this box, and subordinates did not even try to solve it.

While we predicted that vigilance rates would be greater and handling times would be faster when same or higher-ranked audience members were closer to an individual solving a box, the effect of audience distance varied and indeed we saw tendencies in the opposite direction. At the Easy box, same or higher ranked audience members at 20-30 m led to decreased vigilance but no effect on handling times. While at the Medium box, competitors at 20-30 m led to the predicted effects of shorter handling times with greater vigilance. However, competitors at 30-40 m led to longer handling times and greater vigilance. At the Hard box, there were increases in vigilance when competitors were 20-30 m away, but this was not significant and there was no impact of competitor proximity on handling time. One thing we can say confidently from these mixed results is that closer competitors (0-20 m) did not increase vigilance and decrease handling times as we predicted, perhaps because vervets were aware that they were likely to soon be displaced (e.g., Arseneau-Robar et al. 2022) and focused their attention on the box rather than looking around.

We hypothesized that vigilance would be greater for subordinates relative to dominants at every box. While there was an uptick in vigilance with lower dominance for the Easy and Medium box, these trends were not significant (Fig. 5). However, at the Hard box, subordinates were significantly more vigilant than dominants and this was seen even though only Tier 1 and 2 individuals participated. This confirms that even the fairly high ranked individuals in Tier 2 feared displacement at the Hard box by the most dominant individuals while they worked to complete the three-step solving sequence that was required.

If vervets faced a trade-off between attention to the puzzle boxes and competitors, we predicted that they would show slower handling times when they showed greater vigilance rates. This was based on previous literature that proposed that food items that require visual attention while handling should lead to decreases in the time that can be allocated to scanning the surrounding environment (i.e., for predators and conspecifics) (*Parus caeruleus*, Kaby and Lind 2003; *Cercopithecus mitis erythrarchus*, Cowlishaw et al. 2004; four ganivorous bird species, Baker et al. 2011; *Colobus vellerosus*, Teichroeb and Sicotte 2012), and other studies that showed that vigilance to nearby competitors to defend food from kleptoparasitism decreased food intake rates (*Haematopus ostralegus*, Goss-Custard et al. 1999; *Anser alifrons*, Zhao et al. 2020; *Sciurus carolinensis*, Teichroeb et al. 2024). Our results were opposite to our predictions, in that vigilance rates were greater when handling times were faster. We explain this result by invoking the important distinction between scanning frequency and duration. Our measure of vigilance rates quantified head turns per second (scanning frequency), but we did not measure looking duration (scan duration). In samango monkeys (*Cercopithecus mitis erythrarchus*), while scanning duration had a strong negative effect on feeding rates, this effect was less severe for scanning frequencies because quick glances towards threats still allowed some focus on food acquisition (Cowlishaw et al. 2004). Similarly, chacma baboons (*Papio ursinus*) that were in the process of handling challenging foods were found to scan their surroundings more frequently but decrease their duration of looking (Allan et al. 2024). Together, these findings inform our results, which show that vervets that feared displacement at high handling resources could both invest in many quick glances to assess the location of competing conspecifics and speed up their handling time to ensure they were able to access rewards. Each of our puzzle boxes required stepwise solutions that could be completed interspersed with scans, and overall solutions did not require focused attention of long duration. Thus, there is not necessarily a trade-off between attention to foraging tasks that require handling and vigilance if a high scanning frequency to both can be balanced.

Several limitations should be considered when interpreting these findings. The number of individuals completing all three puzzle box conditions was small, reducing statistical power for matched comparisons of first-trial performance. Additionally, dominance interactions were examined using categorical dominance tiers, which may not fully capture the complexity of social relationships within the study group. Future work incorporating continuous dominance ranks and measures of scanning duration to both experimental tasks and competitors would further expand our understanding of these nuanced interactions.

Overall, this study demonstrates that wild vervet monkeys vary in their willingness to engage tasks that require long handling times depending on their dominance. High ranking individuals, that do not fear displacement and aggression, are much more likely to pay handling time costs to gain favoured rewards. We also show that vervets rapidly improve in their problem-solving performance through repeated experience, even when faced with increasingly complex tasks and that they can simultaneously invest in high scanning frequencies to food handling and competitors. These findings contribute to our understanding of how ecological challenges and social environments jointly influence foraging decision-making in wild animals.

## Acknowledgements

We are grateful to Coco Tucker-Gritt, Frances Adams, Dr. Dennis Twinomugisha, and Herman Kasozi for making this research possible.

## Statements and Declarations Funding

This research was funded by a Natural Sciences and Engineering Research Council of Canada Discovery Grant to J. Teichroeb (RGPIN-2023-03613).

## Author Contributions

Ajay S. Kang – Conceptualization, Data curation, Formal analyses, Investigation, Methodology, Project administration, Validation, Visualization, Writing – original draft, Writing – review & editing; Wilson Mutebi – Investigation; Julie A. Teichroeb - Conceptualization, Funding acquisition, Methodology, Project administration, Resources, Supervision, Writing – original draft, Writing – review & editing

## Ethics Approvals

The methods for this experiment were approved by the Uganda Wildlife Authority (Permit: UWA/COD/96/05), the Uganda National Council for Science and Technology (Permit: NS131ES), and the University of Toronto Animal Care Committee (Protocol: 20012354). No animals were captured, handled, or restrained during the study.

## Competing Interests

The authors declare no competing interests.

## Data Availability Statement

The data analyzed in this paper are available from the corresponding author on request.

## Notes

### Competing Interest Statement

The authors have declared no competing interest.

